# Genotyping complex structural variation at the malaria-associated human glycophorin locus using a PCR-based strategy

**DOI:** 10.1101/758334

**Authors:** Walid Algady, Eleanor Weyell, Daria Mateja, André Garcia, David Courtin, Edward J Hollox

## Abstract

Structural variation in the human genome can affect risk of disease. An example is a complex structural variant of the human glycophorin gene cluster, called DUP4, which is associated with a clinically-significant level of protection against severe malaria. The human glycophorin gene cluster harbours at least 23 distinct structural variants and accurate genotyping of this complex structural variation remains a challenge. Here, we use a PCR-based strategy to genotype structural variation at the human glycophorin gene cluster. We validate our approach, based on a triplex paralogue ratio test (PRT) combined with junction-fragment specific PCR, on publically-available samples from the 1000 Genomes project. We then genotype a longitudinal birth cohort using small amounts of DNA at low cost. Our approach readily identifies known deletions and duplications, and can potentially identify novel variants for further analysis. It will allow exploration of genetic variation at the glycophorin locus, and investigation of its relationship with malaria, in large sample sets at minimal cost, using standard molecular biology equipment.

## Introduction

The malaria parasite *Plasmodium falciparum* is a major cause of childhood mortality in Africa (Liu et al. 2016; Snow et al. 2017; Whitty and Ansah 2019). Genome-wide association studies have identified multiple alleles that increase protection from severe malaria symptoms; understanding the mechanistic basis of this protection, the precise phenotype affected by the allelic variation, and the evolutionary history of the protective alleles is an important area of current research (Network 2015; Ravenhall et al. 2018). One of the association signals was shown to be due to a complex structural variant, called DUP4, involving the human glycophorin gene cluster on chromosome 4 (Leffler et al. 2017; Algady et al. 2018).

The human glycophorin genes *GYPE, GYPB* and *GYPA* are located on three paralogous repeat units ∼120kb in size sharing 97% identity at chromosome 4. *GYPA* and *GYPB* encode glycophorin A and glycophorin B respectively, both of which are major components of the erythrocyte membrane and are receptors for ligands on the surface of the *P. falciparum* merozoite (Pasvol et al. 1982; Orlandi et al. 1992; Sim et al. 1994; Mayer et al. 2009; Baldwin et al. 2015). No protein product of *GYPE* has yet been detected, and it is thought therefore to be a pseudogene. There is extensive structural variation at this locus resulting in at least 8 distinct deletions of one or two repeat units, 13 distinct duplications of one or two repeat units, and two more complex variants (Leffler et al. 2017)((Louzada et al. 2019)). These structural variants can be identified and genotyped using sequence read depth of mapped high throughput sequencing reads (Leffler et al. 2017; Algady et al. 2018). However, the ability to rapidly genotype the different structural variants at this locus using simple PCR approaches would have a number of benefits. Firstly it would allow access to large cohorts with limited DNA, where whole genome amplification is both prohibitively costly and known to introduce bias in copy number measurement (Pugh et al. 2008; Veal et al. 2012). Secondly it would allow laboratories in resource-limited situations to genotype glycophorin structural variation, as only standard molecular genetic laboratory equipment, and an instrument suitable for DNA fragment analysis, are needed. Previously we published a PCR based approach to genotype the malaria-protective DUP4 variant using a paralogue specific PCR targeting a junction unique to that variant. We used this PCR approach to genotype a Tanzanian cohort and demonstrate association of the DUP4 variant with hemoglobin levels in peripheral blood (Algady et al. 2018). In this report we describe a simple triplex paralogue ratio test system based on a single PCR of 10ng DNA that can distinguish other structural variants of the glycophorin gene cluster.

The paralogue ratio test (PRT) is a well-established form of quantitative PCR that is particularly robust in copy number detection in genomic DNA. It relies on co-amplification of a test and reference locus using the same PCR primer pair, and distinguishing the two products either by single nucleotide difference (Saldanha et al. 2011) or, more commonly, by a small size difference between the products (Armour et al. 2007). Because the same primer pair is used to amplify test and reference, regions that are similar in sequence, such as segmental duplications or diverged, dispersed, repeats, are usually targeted. The similarity in sequence means that the amplification of both the test and reference loci follow similar kinetics, and quantification of endpoint products reflects the relative amounts of the starting molecules. PRT has been shown empirically to be more robust that more common forms of qPCR that use different primers to amplify test and reference (Aldhous et al. 2010; Fode et al. 2011; Cantsilieris et al. 2014).

We also present a strategy to confirm the most common variants by junction fragment PCRs. Junction fragment PCR relies on knowledge of the copy number breakpoint to design paralogue-specific allele-specific primers flanking the breakpoint, so that if the copy number variant is present, the two paralogue-specific primers are brought into close proximity, enabling a short PCR product to be successfully amplified. Using these approaches, we present a cost-effective genotyping strategy structural variation in the human glycophorin region to facilitate large-scale genotyping studies of valuable DNA cohorts.

## Methods

### DNA samples

Control DNA samples from the 1000 Genomes project were chosen because DNA samples are publically available, and high throughput Illumina sequencing has been used to genotype structural variation at this locus (Sudmant et al. 2015; Leffler et al. 2017). A subset of genotypes of this sample set have been validated by fibre-FISH and breakpoints identified ((Louzada et al. 2019)). DNA samples for CEPH Europeans from Utah (CEU), Chinese from Beijing, Japanese from Tokyo and Yoruba from Ibadan Nigeria (YRI) were previously purchased from Coriell Cell Repositories. DNA from the Beninese infants came from a longitudinal birth cohort established in the district of Tori Bossito (Le Port et al. 2012). The protocol was approved by both the Ethical Committee of the Faculté des Sciences de la Santé (FSS) in Benin and the IRD Consultative Committee on Professional Conduct and Ethics (CCDE). All genomic DNA samples were diluted to a concentration of 10ng/µl in water

### Paralogue ratio test

Triplex PRT was conducted essentially as previously published (Armour et al. 2007; Walker et al. 2009; Hollox 2017). Briefly, three primer pairs targeting different regions of the glycophorin repeat were combined in a single PCR reaction as follows: In a final volume of 10µl, 10ng of genomic DNA was amplified using 0.5µl each of 10µM forward and reverse primers for the three PRT assays (table 1), 1µl of 10x low dNTP buffer (Hollox 2017), 1µl of 10x *Taq* DNA polymerase buffer (KAPA Biosystems, with 15mM MgCl2), 0.5U *Taq* DNA polymerase. The forward primer of each PRT assay was 5’ labelled with a fluorescent tag, either 6-FAM or HEX, to allow visualisation of products by capillary electrophoresis. Cycling conditions were an initial denaturation at 94°C for 2 minutes, followed by 25 cycles of 94°C 30 seconds, 65°C 30 seconds, 70°C 30 seconds, and a final extension of 70°C for 5 minutes. PCR products were denatured and run on an ABI3130xl capillary electrophoresis machine using POP-4 polymer with an injection time of 30s under standard fragment analysis parameters and ROX-labelled Mapmarker 1000XL (Eurogentec) and areas under the peak of each PCR product subsequently analysed using the GeneMapper software. Samples with PCR product peak areas between 300 and 60000 were accepted.

**Table 1.**
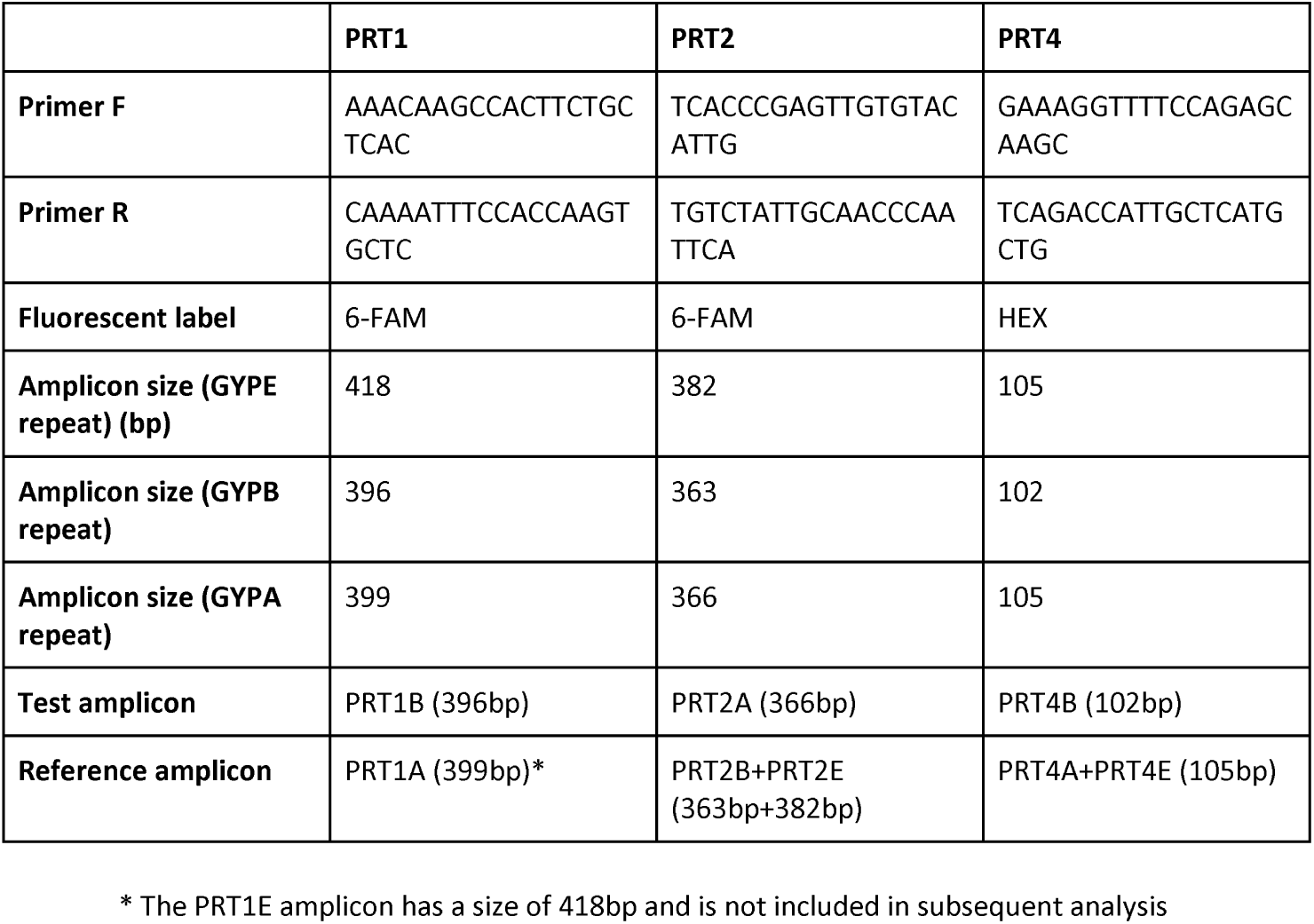
PRT primers and product details.

For each sample, PRT area ratios using test and reference peaks as table 2 and normalised using DNA samples of known copy number ((Handsaker et al. 2015; Leffler et al. 2017) (Louzada et al. 2019)) that were included in every experiment: NA19190 (CN=1, DEL2), NA19818 (CN=1, DEL2), NA19085 (CN=2, reference), NA19777 (CN=2, reference), NA19084 (CN=3, DUP2). Following normalisation, data were further normalised to account for small batch effects by dividing against the median value for that experiment.

**Table 2.**
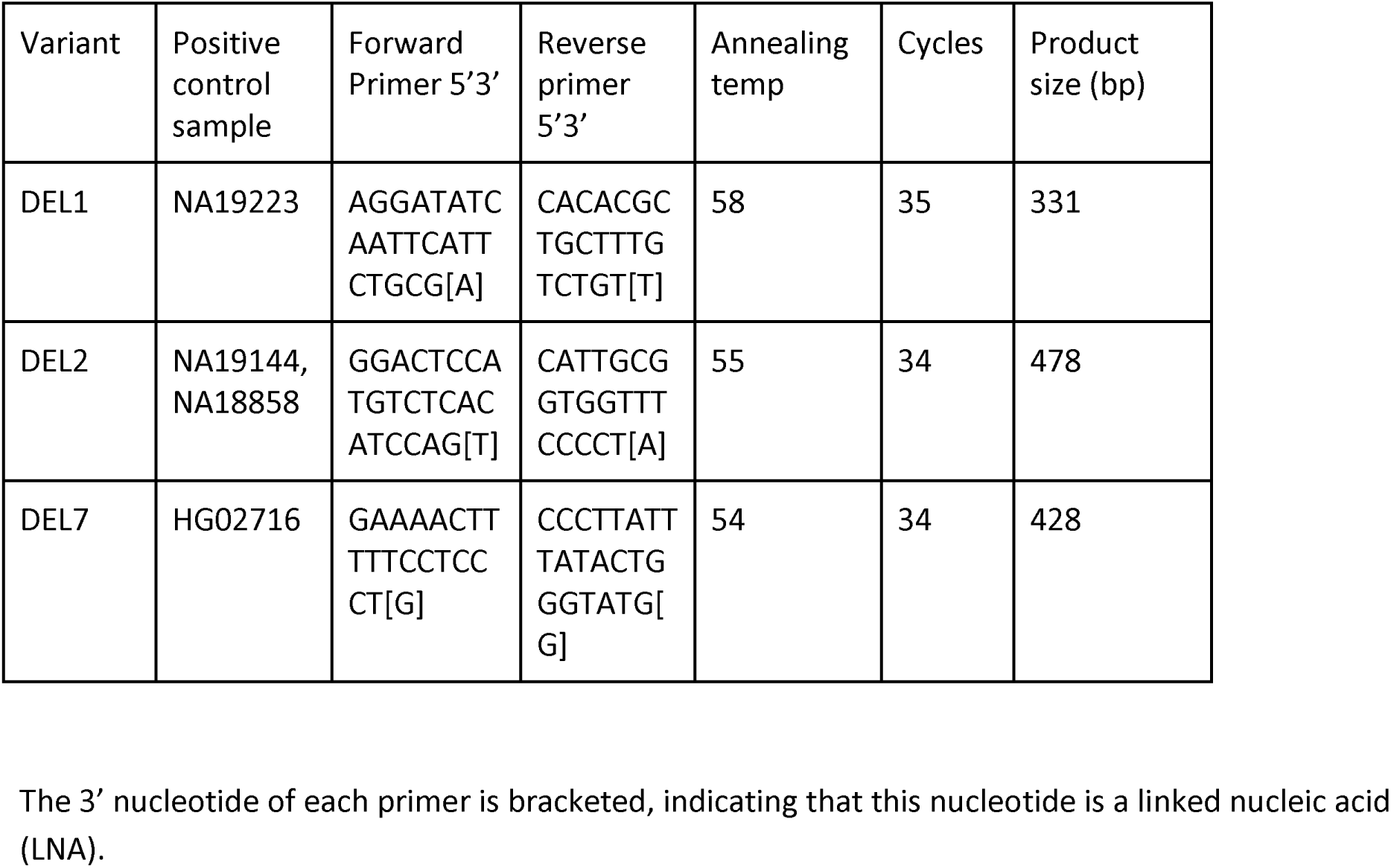
Junction fragment PCR primers and product details.

### Junction fragment duplex PCR

Junction fragment duplex PCR was conducted using 10ng genomic DNA, 10pmol of each breakpoint-specific primer with a 3’terminal linked nucleic acid to enhance specificity (table 2), 1pmol of each control primer (5’GAGTACGTCCGCTTCACC-3’ and 5’CTTCCACACTTTTGGCATGA-3’), 0.5U *Taq* DNA polymerase, 2µmol each of dATP, dCTP, dGTP, dTTP in a final volume of 10µl 1x Kapa Buffer A (ammonium sulfate-based buffer, final concentration MgCl2 1.5mM). PCR cycling was 95°C for 2 minutes, followed by x cycles of 95°C 30s, y°C, 30s, 72°C, 30 seconds with a final extension of 72°C for 5 minutes. The number of cycles (x) and annealing temperature (y) were assay-specific, and shown in table 2. PCR products were analysed using standard 2% agarose gel electrophoresis, staining by ethidium bromide and visualisation by UV light. The control primers amplify a 216bp product, and the breakpoint-specific primers amplify product sizes shown in table 2.

## Results

### Development of triplex PRT system

A triplex PRT consists of three distinct PRTs that are amplified together in a single PCR reaction, which generates three independent measures of copy number in one single tube and electrophoresis capillary (Walker et al. 2009). If the PRTs have a test and reference on the same chromosome, they are called cis-PRTs, in contrast to trans-PRTs where the reference is located on a different chromosome to the test amplicon (Hollox 2017). In this study we designed three cis-PRTs, each one targeting a different region of the glycophorin repeat, with test and reference distinguished on the bases of product size by electrophoresis (figure 1). Because the most frequent deletion affects the GYPB gene, the test amplicon was the GYPB amplicon with the reference amplicons being either GYPA or GYPE amplicons. Since the extent of the most frequent structural variants are known, and the three different PRTs measure copy number at different regions of the glycophorin repeat, it is possible to predict the relative test/reference ratios of these different structural variants (figure 2).

**Figure 1.**
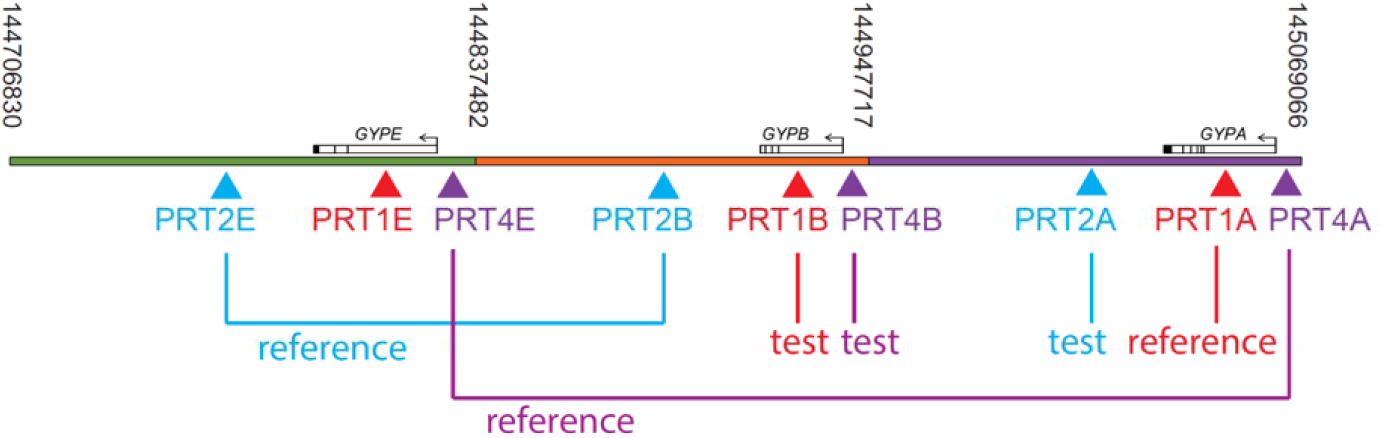
Multiplex paralogue ratio test approach in measuring structural variation at the human glycophorin gene cluster. The hg19 Grch37 reference sequence is shown, with numbers indicating boundaries between the three glycophorin repeats coloured green, orange and purple carrying GYPE, GYPB and GYPA respectively. Amplicons for PRT1, PRT2 and PRT4 are indicated by triangles. Test and reference amplicons for each PRT are also shown.

**Figure 2.**
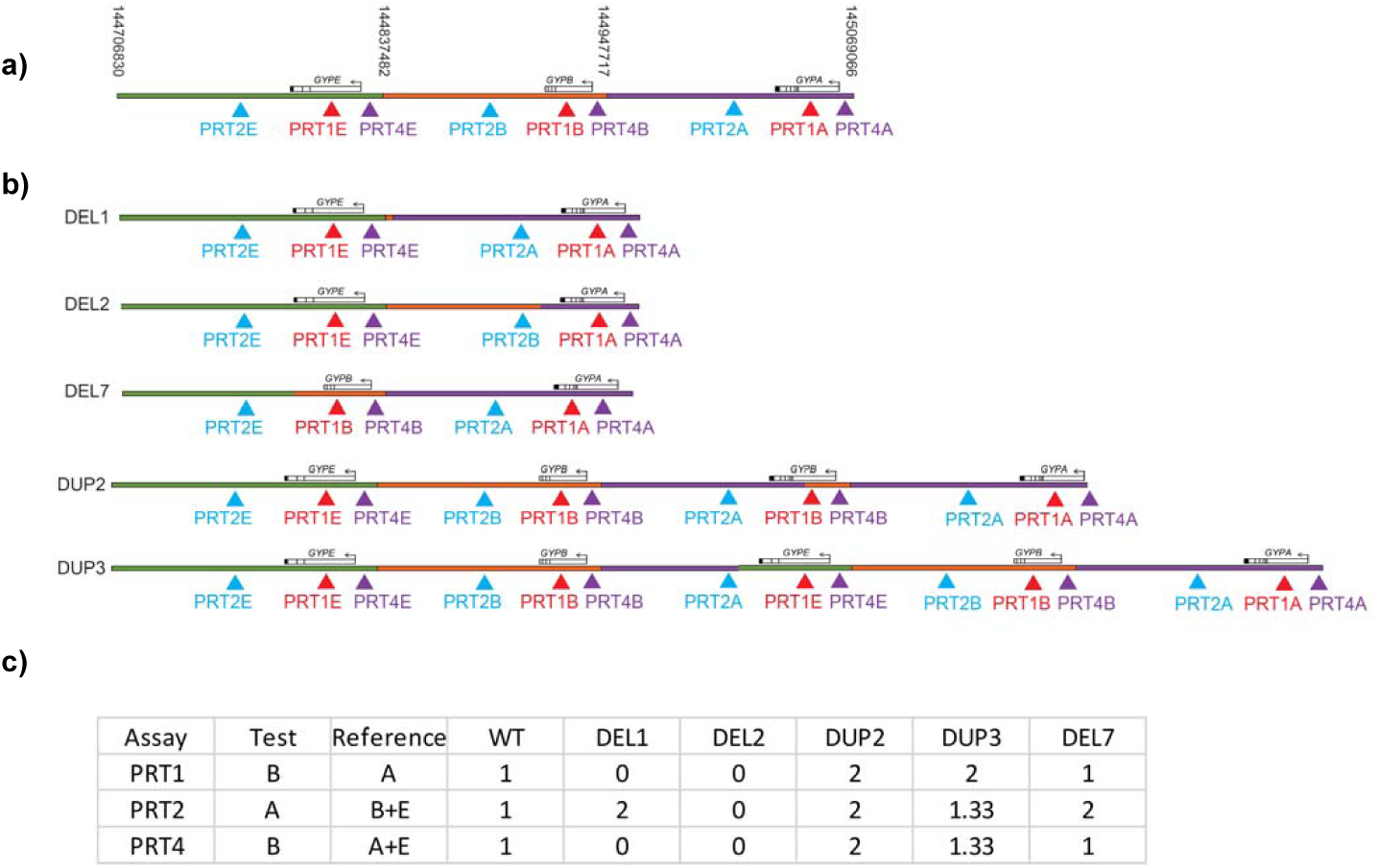
Distinguishing different structural variants using the multiplex paralogue ratio test. a) The hg19 GRCh37 reference genome, together with locations of PRT amplicons. b) Structures of five variants (DEL1, DEL2, DEL7, DUP2 and DUP3) that have been determined previously by fibre-FISH are shown, together with PRT amplicon sites. c) Expected dosage values for each PRT assay for the reference genome (wild-type, WT) and the five variants shown. Note that the PRT amplicons used as test and reference are shown, and the reference wild-type variant defined with a value of 1. For example, the ratio of PRT1A to PRT1B is 1 for the WT allele, and therefore predicted to be 2 for DUP3 (two PRT1B sites, one PRT1A site).

We validated our triplex PCR by analysing a cohort of 177 1000 Genomes samples with matching short read sequence data, where structural variants of the glycophorin region previously identified. We could then directly compare our PRT results with estimates of copy number from previous sequence read depth analysis and fiber-FISH ((Louzada et al. 2019)). Analysis of PRT1 shows complete concordance with NGS-measured copy number, detecting the deletions and duplications identified previously (figure 3a). Analysis of PRT2 shows a more complex situation; although the duplications are identified correctly, some of the deletion samples are identified by a gain of signal by PRT2 rather than a loss of signal (Figure 3b). This discrepancy is expected, as PRT2 measures DEL2 as an increase in signal, because the amplicon corresponding to GYPA (reference) is lost not GYPB (test) (figure 2). Three outliers were not previously identified as known structural variants and are likely to represent a variant with a gene conversion event affecting one of the PRT amplicons (Figure 3b). PRT4 yields similar results to PRT1 for this cohort, with a few individuals with no structural variant identified showing a higher than expected value, again probably due to gene conversion events.

**Figure 3.**
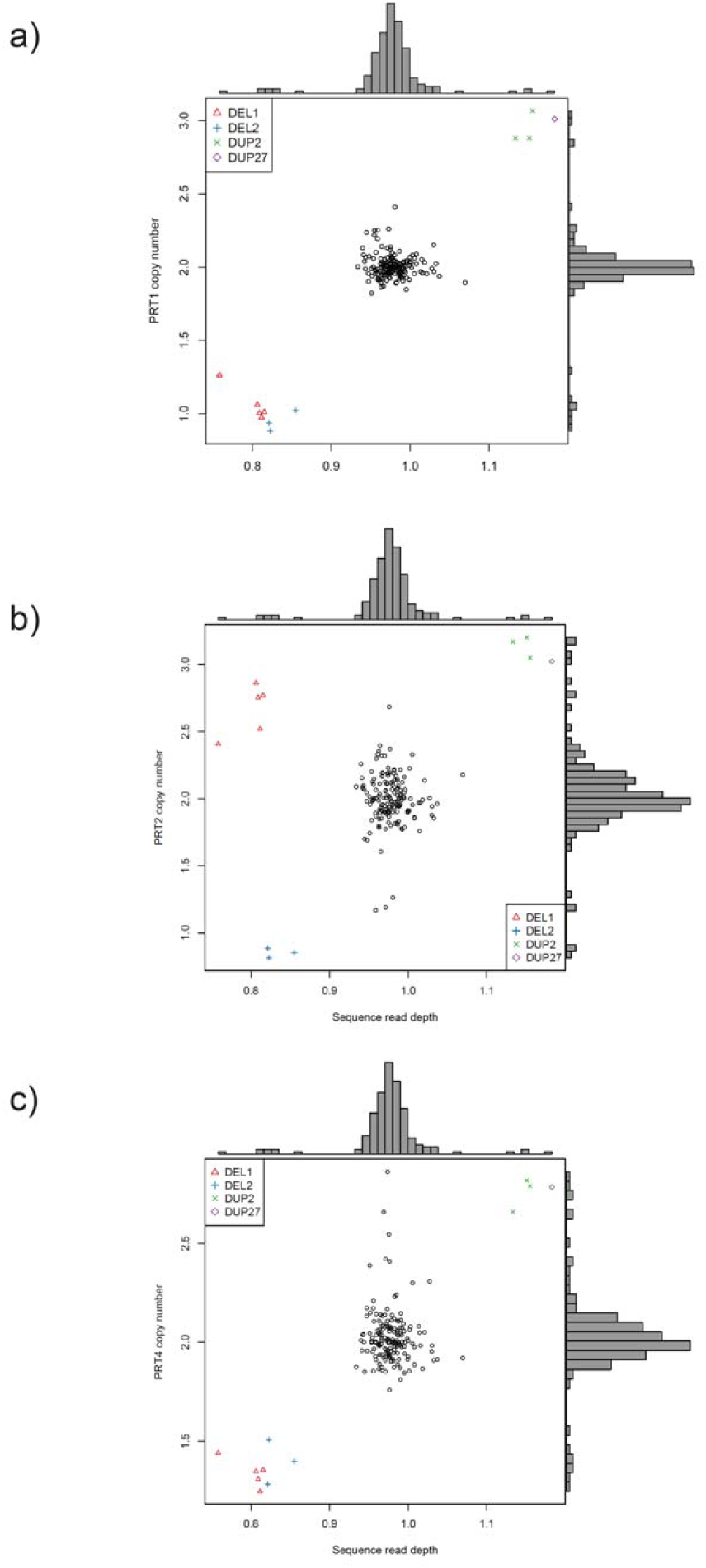
Comparison of PRT-based copy numbers to sequence read depth-based copy numbers in 1000 Genomes samples. The scatterplots show comparison of sequence read depth estimation of copy number (with 1.0 being normal read depth) on the x axis and PRT copy number estimates on the y axis for a) PRT1, b) PRT2 and c) PRT3. Each point represents a different individual, with point shape/colour indicating the inferred structural variant carried as a heterozygote by that sample, as shown in the legend. Black circles indicate samples interpreted as homozygous wild-type reference variants.

By comparing across all three PRTs, the four different structural variants can be distinguished from individuals not carrying a structural variant. The exception is distinguishing DUP2 and DUP27, which are either the same variant, or share very close breakpoints ((Louzada et al. 2019)). This analysis shows that by using different loci as test and reference loci, and by combining PRT assays together, different structural variants can be distinguished on the basis of copy number and breakpoint (table 3). It should also be noted that, for PRT4, the clusters for the deletions are higher than expected, with a median value for heterozygotes around 1.25 compared to a theoretical value of 1 (table 3), and a median value for homozygotes of 0.5 compared to a theoretical value of 0 (table 3). Of the two DEL1 control samples, NA19818 shows the expected level of test amplicon amplification while NA19190 shows a much lower level. The reason for this is unclear – it might be due to an undetected variant in the PRT4 test locus primer binding site on the reference allele in NA19190 – but the consequence is that the calibration curve for the positive controls for PRT4 is slightly distorted at lower copy number values. For future studies, we recommend using only NA19818 as the DEL1 positive control.

**Table 3.**
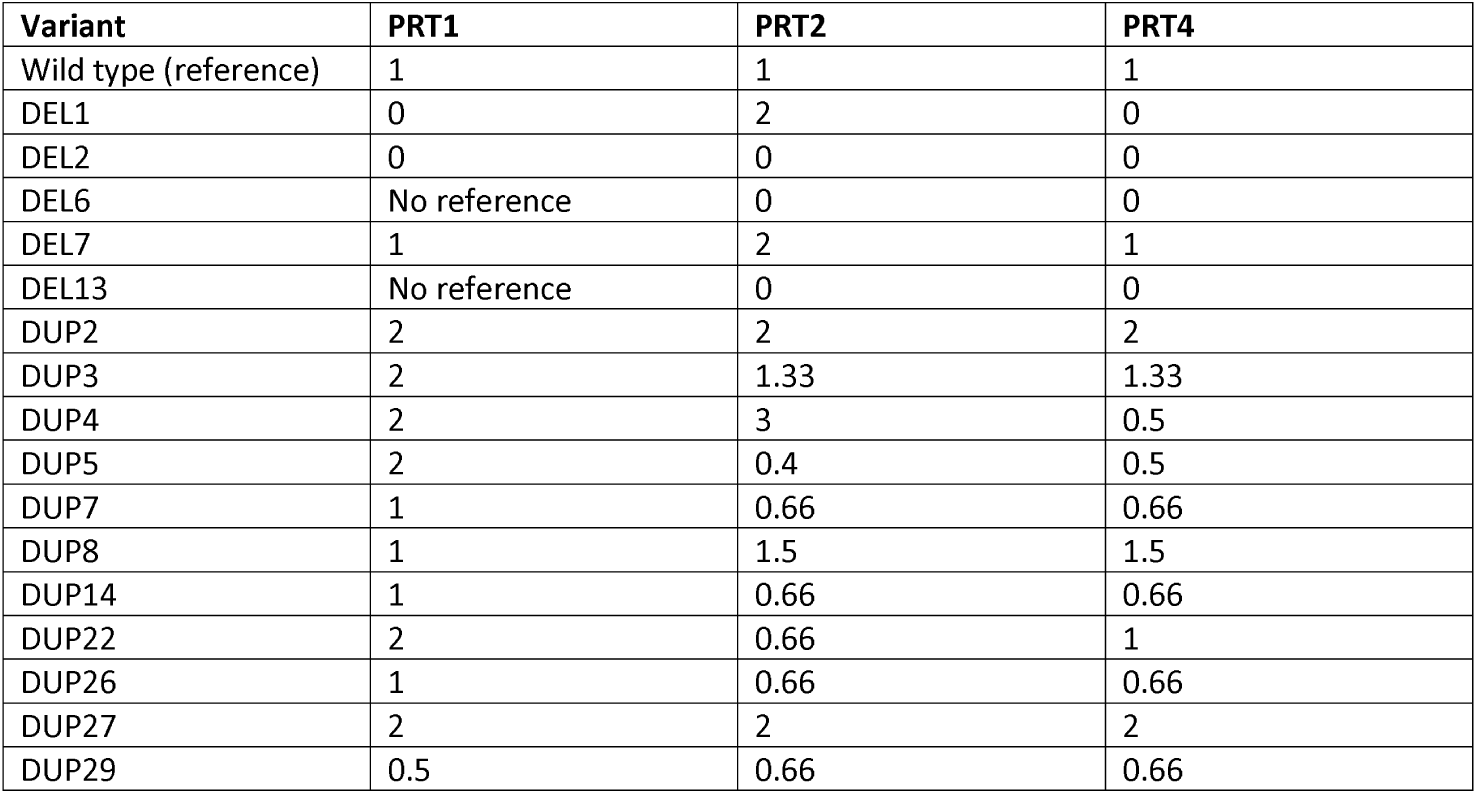
Predicted normalised PRT ratios for glycophorin variants observed in 1000 Genomes project cohort.

### Analysis of Benin population

We used our triplex PRT approach on a cohort of 574 individuals from Benin in west Africa. Individuals from west African populations have been shown to have appreciable frequencies of different glycophorin structural variants (Leffler et al. 2017). A plot of PRT1 copy number estimates against PRT2 copy number estimates shows clear clustering of individuals (Figure 4a), with genotype inferred using the relative dosages predicted from the structures of the different variants (Figure 2). A plot of PRT1 copy number estimates against PRT4 copy number shows dosages consistent with the assigned genotypes, but with several WT/WT samples as outliers for PRT4. The molecular basis for these outliers is not yet clear, they have not previously been identified as novel structural variants affecting copy number, so it seems likely that they are due to gene conversion events altering the ratio of test/reference amplicon dosage without altering the overall copy number of the locus.

**Figure 4.**
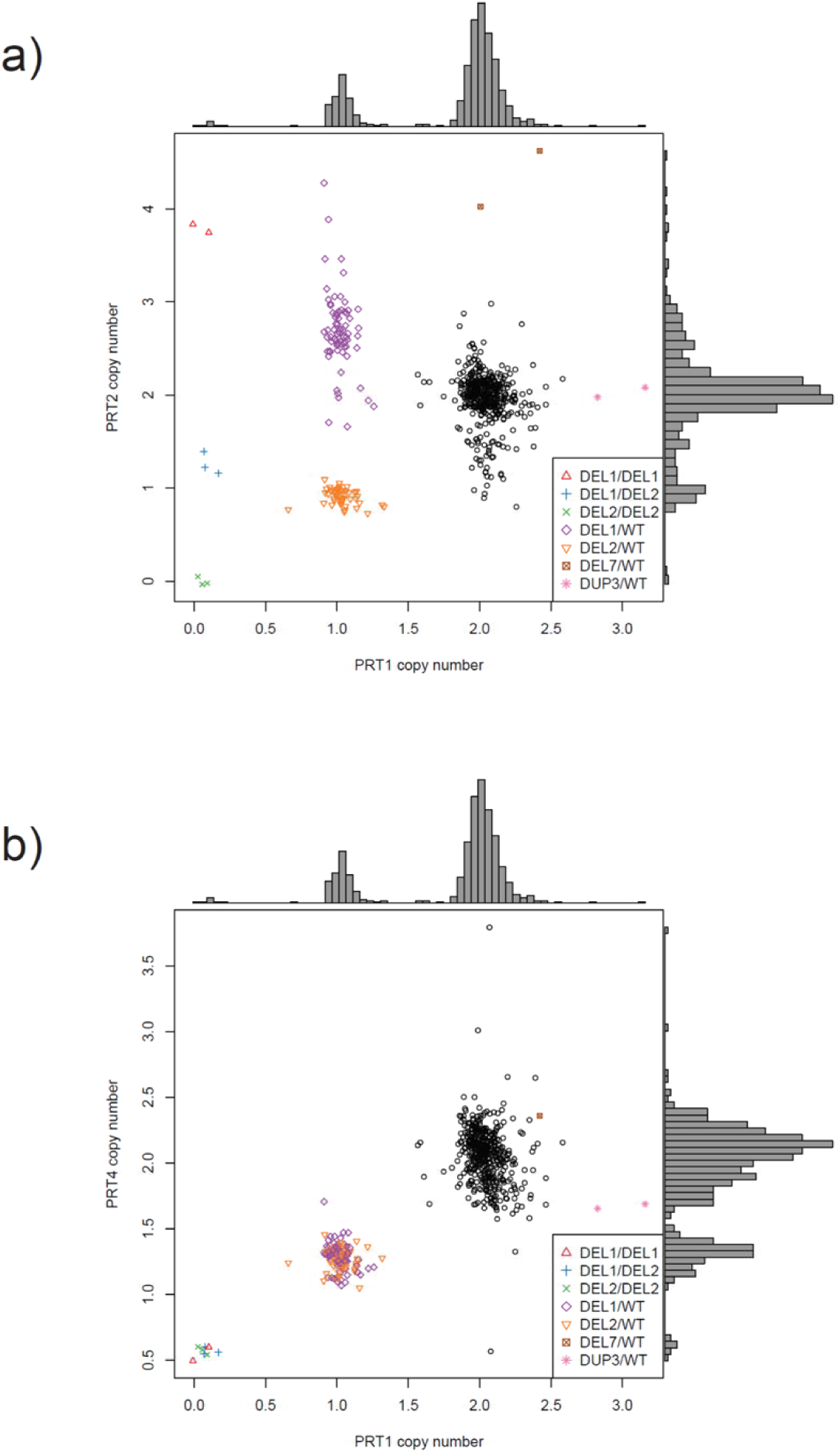
Comparison of copy numbers from different PRT tests allows determination of structural variant genotype. The scatterplots show comparison of PRT copy number estimates for a) PRT1 versus PRT2 and b) PRT1 versus PRT4 on a cohort of individuals from Benin. Each point represents a different individual, with point shape/colour indicating the inferred structural variant genotype. Black circles indicate samples interpreted as homozygous wild-type reference variants.

### Confirmation using junction-fragment

*PCR* To confirm genotypes, we designed junction-fragment PCRs to span the known breakpoints of DEL1, DEL2, as determined previously (Louzada et al. 2019). These were designed to generate short amplicons with a co-amplified non-variable control amplicon from an unrelated locus in the genome, to act as a positive control for DNA amplification. To ensure paralogue-specificity, the 3’ end of each primer was designed over a nucleotide that distinguished the paralogue to be amplified from the other two paralogues, and was synthesised using a linked nucleic acid nucleotide analogue to enhance specificity in base-pairing (Latorra et al. 2003).

## Discussion

In this study we present a genotyping strategy for structural variants of the glycophorin gene cluster in humans. We use a PCR-based approach called the paralogue ratio test (PRT) to estimate copy number gain and loss across the region. By combining information from three separate PRT assays, the genotype of particular structural variants can be called. These can then be confirmed by paralogue-specific junction-fragment PCR assays.

There are several strengths in the PRT approach. It calls copy number and does not rely on homogeneity of variant breakpoints, unlike junction fragment PCR. This allows a variant to be called that may not share the exact breakpoint with other examples of the same apparent variant, and this is known to be a complication in DUP2, encoding the Sta antigen. Because PRT is based on PCR of small fragments, it can be applied to large sample cohorts containing small quantities of degraded DNA. The two-stage approach of PRT followed by junction-fragment PCR allows initial triage of samples homozygous for the reference variant and selection of samples carrying rarer variants for confirmatory junction-fragment PCR, or other analyses, such as targeted high-throughput sequencing. Our approach uses widely-available low-cost molecular biology equipment, with the exception of capillary electrophoresis equipment. Our chosen positive controls are form the 1000 Genomes project, which have been validated extensively by short-read sequence read depth analysis, fibre-FISH and breakpoint Sanger sequencing ((Louzada et al. 2019)). It is also adaptable, for example using PRT1 alone would be the most straightforward to interpret for common deletions and duplications, and can be used to select for particular genotypes of interest for sequencing, for example.

We demonstrated the usefulness of our approach by genotyping 574 individuals from a Beninese birth cohort. We found two deletions also observed in other west African populations (DEL1 allele frequency 7%, DEL2 allele frequency 5%), with genotype frequencies in Hardy-Weinberg proportions, but there is no previous evidence that these affect the risk of malarial clinical phenotypes (Leffler et al. 2017). We found two examples of the DEL7 variant and two DUP3 variants, confirming that these as present, but rare, in west African populations. We found no examples of the DUP4 variant that is protective against malaria.

As with any approach, there are weaknesses. As mentioned above, the junction PCR assumes homogeneity of breakpoints across all copies of the same variant. While PRT can determine copy number robustly (Cantsilieris et al. 2014; Adewoye et al. 2018), care must be taken to include internal positive controls to allow for normalization within an experiment and across experiments (Hollox 2017).

In conclusion, we hope our genotyping approach will be useful to other investigators in tackling this fascinating, complex region where variation has profound consequences on the susceptibility of individuals to malarial disease.

## Acknowledgements

We would like to thank Rachael Madison for technical support, and Mark Jobling for access to the ABI 3100xl capillary electrophoresis machine. We would also like to thank the children and families from Tori Bossito in Benin for their contribution to this study.

